# Network-based asymmetry of the human auditory system

**DOI:** 10.1101/251827

**Authors:** Bratislav Mišić, Richard F. Betzel, Alessandra Griffa, Marcel A. de Reus, Ye He, Xi-Nian Zuo, Martijn P. van den Heuvel, Patric Hagmann, Olaf Sporns, Robert J. Zatorre

## Abstract

Converging evidence from activation, connectivity and stimulation studies suggests that auditory brain networks are lateralized. Here we show that these findings can be at least partly explained by the asymmetric network embedding of the primary auditory cortices. Using diffusion-weighted imaging in three independent datasets, we investigate the propensity for left and right auditory cortex to communicate with other brain areas by quantifying the centrality of the auditory network across a spectrum of communication mechanisms, from shortest path communication to diffusive spreading. Across all datasets, we find that the right auditory cortex is better integrated in the connectome, facilitating more efficient communication with other areas, with much of the asymmetry driven by differences in communication pathways to the opposite hemisphere. Critically, the primacy of the right auditory cortex emerges only when communication is conceptualized as a diffusive process, taking advantage of more than just the topologically shortest paths in the network. Altogether, these results highlight how the network configuration and embedding of a particular region may contribute to its functional lateralization.

## INTRODUCTION

The brain is a complex network of anatomically connected and functionally interacting neuronal populations. These connectivity patterns span multiple spatial and topological scales [6, 40], conferring the capacity for both specialized processing and multimodal integration among distributed systems. Increasing evidence suggests that the anatomical connectivity patterns may not be perfectly symmetric however, with several systems marked by lateralized connection density and topological features [36].

Auditory networks in particular display a pronounced tendency for functional asymmetry [11]. Numerous studies have reported both structural and functional differences between the left and right auditory cortex, and have documented their differential contributions to a wide range of sensory and cognitive tasks, including speech [27] and tonal processing [67]. These asymmetries have also been observed at the network level, with asymmetric patterns of functional interactions or connectivity during specific tasks involving auditory processing, such as speech and language [48] and pitch processing [13].

Recent evidence from stimulation studies raises the possibility that this lateralization is mediated by asymmetric anatomical connectivity and network embedding of the auditory cortices. For instance, stimulation of the auditory network by transcranial magnetic stimulation (TMS) elicits highly asymmetric patterns of activity and functional connectivity, with more widespread effects if the stimulus is applied over the right auditory cortex compared to the left [2]. Importantly, individual differences in responses to stimulation are predicted by both interhemispheric anatomical connectivity and resting state functional connectivity [1]. Altogether, these studies suggest that the functional asymmetry of the auditory network may partly be a consequence of its topology, with the right auditory cortex better positioned to disseminate, exchange and integrate neural signals with other systems.

Here we investigate whether the observed functional asymmetry of the auditory system can be attributed to the anatomical network embedding of left and right auditory cortex. Using connectivity patterns reconstructed from diffusion weighted imaging in three different datasets, we assess the propensity for left and right auditory cortex to maintain connections and potential communication pathways with the rest of the brain.

Importantly, we investigate a range of measures that embody different models of network communication. To assess the potential for left and right auditory cortex to communicate with the rest of the brain via shortest paths, we estimate the path length between these areas and the rest of the network (also referred to as closeness centrality or nodal efficiency). To assess the potential for these areas to communicate via an ensemble of paths, we estimate their communicability with the rest of the network [3, 17, 20, 24, 59]. Finally, we use a simple spreading model in which focal perturbations in left and right auditory cortex develop into global signaling cascades that diffuse through the network [31, 45, 61, 63]. Unlike path length and communicability, the model is inherently dynamic, and allows us to trace the trajectories of putative signaling cascades. We hypothesize that if the lateralization of the auditory system has an anatomical origin, the network embedding of the primary auditory cortices will differ between the left and right hemispheres, with the right auditory cortex better positioned to communicate with, and influence, other areas.

## METHODS

### Datasets

We performed all analyses in three diffusion-weighted imaging datasets. The main (discovery) dataset was collected at the Department of Radiology, University Hospital Center and University of Lausanne, (LAU; N=40). We also included two replication cohorts, one from the Human Connectome Project (HCP; N=215) [60] and Nathan Kline Institute Rockland Sample (NKI; N=285)

[49]. Structural connectivity was reconstructed from diffusion-weighted imaging: diffusion spectrum imaging (DSI) for LAU, high angular resolution diffusion imaging (HARDI) for HCP and diffusion tensor imaging (DTI) for NKI. Although dataset LAU had the fewest participants, we selected it as the main dataset to demonstrate our findings because of the quality of the DSI sequence. Below we describe the acquisition, processing and connectome reconstruction procedure for each dataset in more detail.

#### LAU

A total of N = 40 healthy young adults (16 females, 25.3 *±* 4.9 years old) were scanned at the Department of Radiology, University Hospital Center and University of Lausanne. Grey matter was parcellated according to the Desikan-Killiany atlas [22]. These regions of interest were further divided into 114 approximately equally-sized nodes [10]. Structural connectivity was estimated for individual participants using deterministic streamline tractography as implemented in the Connectome Mapping Toolkit [10], initiating 32 streamline propagations per diffusion direction for each white matter voxel. For more details regarding the acquisition protocol and reconstruction procedure, see [45].

#### HCP

A total of N = 215 healthy young adults (112 females, 29.7 *±* 3.4 years old) were scanned as part of the HCP Q3 release (Van Essen et al., 2013). Grey matter was parcellated according to the Desikan-Killiany atlas [22]. These regions of interest were further divided into 219 approximately equally-sized nodes [10]. Structural connectivity was estimated for individual participants using generalized q-sampling (GQI) [64] and deterministic streamline tractography. For more details regarding the acquisition protocol and reconstruction procedure, see [44].

#### NKI

A total of N = 285 healthy adults (112 females, 44.38 *±* 19.7 years old) were scanned as part of the NKI initiative [49]. Grey matter was parcellated into 148 regions of interest according to the Destrieux atlas [23]. Structural connectivity was estimated for individual participants using the Connectome Computation System (CCS: http://lfcd.psych.ac.cn/ccs.html). For more details regarding the acquisition protocol and reconstruction procedure, see [6].

### Defining auditory and visual regions

Primary auditory and visual cortex were delineated according to the Desikan-Killiany (for LAU and HCP; [22]) and Destrieux atlases (for NKI; [23]). Both atlases are based on automated anatomical labeling of MR images using gyrual and sulcal landmarks. Primary auditory cortex was defined as the “transverse temporal” (Desikan-Killiany) and the “G_temp_up-G_T_transv” (Destrieux) nodes. Primary visual cortex was defined as the “pericalcarine” (Desikan-Killiany) and “S_calcarine” (Destrieux) nodes. None of these nodes were subdivided into smaller units than defined in the original atlases.

### Consensus adjacency matrices

Given recent reports of inconsistencies in reconstruction of individual participant connectomes [37, 57], as well as the sensitive dependence of network measures on false positives and false negatives [65], we adopted a group-consensus approach, whereby for each dataset we estimated edges that occur most consistently across participants [19, 54]. In constructing a consensus adjacency matrix, we sought to preserve (a) the density and (b) the edge length distribution of the individual participants’ matrices [6, 45]. The approach is conceptually similar to the procedures proposed by [19, 54].

We first collated the extant edges in the individual participant matrices and binned them according to length. The number of bins was determined heuristically, as the square root of the mean binary density across participants. The most frequently occurring edges were then selected for each bin. Thus, if the mean number of edges across participants in a particular bin is equal to *k*, we selected the *k* edges of that length that occur most frequently across participants. To ensure that inter-hemispheric edges are not under-represented, we carried out this procedure separately for inter‐ and intrahemispheric edges. The binary densities for the final whole-brain matrices were 20.1% (LAU), 8.2% (HCP) and 11.1% (NKI). For each of the matrices, the densities of the left and right hemispheres were 32.2 vs 32.0% (LAU), 14.3 vs 13.7% (HCP) and 18.8 vs 21.0% (NKI) (Fig. S2).

### Communicability

Communicability (*C_ij_*) between two nodes i and j is a weighted sum of all paths and walks between those nodes [24]. For a binary adjacency matrix A, communicability is defined as

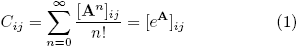

with walks of length *n* normalized by *n*!, such that shorter, more direct walks contribute more than longer walks.

### Linear threshold model

The linear threshold model (LTM) is a simple contagion model that describes how a perturbation introduced at one or more seed nodes develops into a cascade and spreads through a network [31, 61]. The perturbation and subsequent cascade are modeled as an active state; any given node adopts this active state only if a supra-threshold proportion of its neighbors have also adopted the active state. A form of contact percolation, the cascading behaviour described by LTM have been extensively studied over a wide range of networks, including spatially embedded brain networks [38, 39, 45, 50, 63]. The models capture how generic focal perturbations, such as the transduction of a sensory stimulus, spread through connected neuronal populations (see *Discussion* for a discussion of the neurobiological interpretation and limitations).

Formally, the state of a node *i* at time *t* is denoted as a binary variable *r_i_*(*t*) = {0, 1}, with only two possible states: active (1) or inactive (0). At initialization (*t* = 0), the entire network is inactive, except for a subset of activated seed nodes. The model is then updated synchronously at each time step according to the rule:

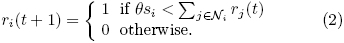

Thus, at each time step the state of node *i* depends on its neighborhood, *N_i_* and specifically on the number of incident connections (degree or strength, *s_i_*). The node adopts the active state only if the proportion of inputs from active nodes exceeds the threshold *θ*. In the case of binary networks, the threshold represents the proportion of a nodeâĂŹs neighbors that must be active to propagate the cascade. The model can be naturally extended to weighted and directed networks, whereby the threshold represents the proportion of a node’s total weighted inputs (strength) that must be connected to active neighbors. In all scenarios, the fundamental performance measure is the adoption or spread time *A_i→k_*, from seed node *i* to target node *k*.

How does the threshold influence spreading dynamics? At lower thresholds, nodes require fewer neighbors to be active at time *t* to become active at time *t* + 1. Thus, nodes will be activated at the earliest possible time step, and the cascade will effectively propagate along the shortest path. As the threshold is increased beyond the inverse of the highest degree/strength in the network, cascades can no longer influence the most highly connected nodes and do not spread through the whole network (Fig. S3). Specifically, at higher thresholds it is more difficult to activate nodes, as more of their neighbours need to be active, so the dynamics are more dependent on local connectivity. At lower thresholds, the dynamics are less constrained by local connectivity and more influenced by global topology.

In the present study, we selected the threshold using the following criteria. The threshold had to be low enough to ensure that all perturbations will cause a complete cascade, so that spread times from the left and right auditory cortex could be unambiguously compared (Fig. S3). Increasing the threshold biases spreading away from shortest paths, with much of the spreading process occurring via alternative paths as well. As a result, spread times become less correlated with path length at greater thresholds (Fig. S4a). We therefore selected a threshold (*θ* = 0.05) at which cascades could reach the whole network. In all three datasets, this corresponded to *θ* = 0.05.

How sensitive is the main effect of interest – the difference in spread time for perturbations originating in left and right auditory cortices – to this parameter setting? Fig. S4b shows the effect of varying the threshold on the left-right auditory cortex asymmetry. At lower thresholds spreading is similar to shortest path routing, and there are no significant differences between left and right auditory cortex. As the threshold is increased, there is a range in parameter space ([0.04 0.09]) where spreading is significantly faster from the right auditory cortex compared to left auditory cortex.

## RESULTS

White matter networks (connectomes) were reconstructed from diffusion weighted imaging (DWI) in three cohorts of healthy adults. We investigated the lateralization of primary auditory cortex by quantifying the topological distance from the left and right auditory cortex to the rest of the brain. We estimated topological distance using three measures, each of which makes different assumptions about the nature of inter-regional communication: path length (the minimum number of edges between two nodes), communicability (weighted sum of all walks between two nodes) and spread time (the time required for a signaling cascade to spread from one node to another; see *Methods* for details of model implementation). The spread time is a dynamic measure of inter-regional communication, estimated by simulating how a focal perturbation develops into a global signaling cascade and spreads through the network.

### Right auditory cortex is topologically more central

To assess the statistical reliability of differences in the anatomical centrality or embedding between the left and right auditory cortex, we used nonparametric tests. In the case of communicability, we used Wilcoxon signed-rank tests [62]. In the case of path length and spread times, which are not continuous, we used unpaired permutation tests (10,000 repetitions).

Two salient findings emerge. First, there are differences in the anatomical embedding of left and right auditory cortex, but these differences only emerge when one considers communication metrics that assume diffusion of information, rather than shortest path routing. Namely, we find that the left and right auditory cortex are indistinguishable in terms of their path length to the rest of the brain (*P* = 0.39). Conversely, the right auditory cortex is topologically closer to other brain areas in terms of diffusive spreading, including greater communicability (*P* = 0.04) and faster spreading times (*P <* 10*^−^*^5^) (Fig. 1, top row). As shown in Fig. S2, the two hemispheres are comparable in their connection density, so the observed effects are more likely to have arisen from differences in topology.

**FIG. 1.**
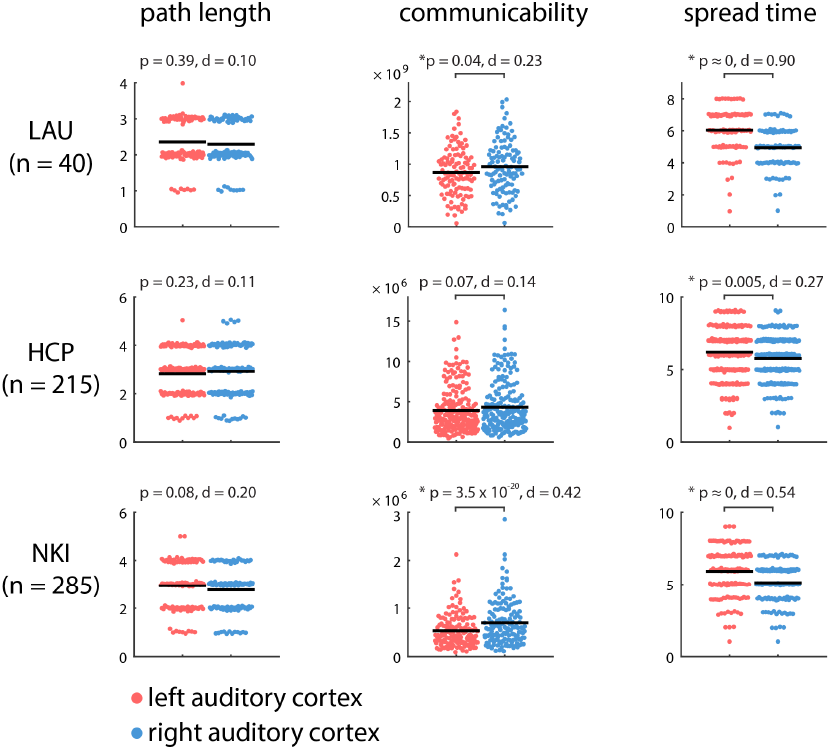
Communication distance from auditory cortices to the rest of the brain. The centrality of left and right auditory cortices was estimated by their topological distance to other brain areas in terms of path length, communicability and spread time. Shorter path length, greater communicability and shorter spread times indicate greater proximity. Mean values for each distribution are indicated by solid horizontal black lines. For visualization, a random horizontal jitter was added to all points. In the case of path length, which is a discrete-valued variable, an additional vertical jitter was added to all points.

Second, despite significant differences in acquisition protocol, processing parameters, resolution and tractography algorithm, these results were replicated in the HCP and NKI datasets (Fig. 1, middle and bottom rows). In both datasets, left and right auditory cortex were statistically indistinguishable in terms of their path length to the rest of the network (*P* = 0.23 in HCP; *P* = 0.08 in NKI), while the spread time for cascades originating in right auditory cortex was significantly faster compared to those originating in left auditory cortex (*P* = 5*×*10*^−^*^3^ for HCP; *P <* 10*^−^*^5^ for NKI). The asymmetry was not only statistically significant, but also associated with a large overall effect size in all three datasets (Cohen’s *d* = 0.90, 0.27, 0.54 for LAU, HCP and NKI datasets, respectively).

### Auditory asymmetries are cumulative

We next sought to pinpoint the origin of these anatomical asymmetries. Are left-right topological asymmetries due to a specific anatomical connection, or do they reflect a more global, cumulative effect? To answer this question, we used the spreading model because (a) it is dynamic, allowing us to trace the evolution of each signaling cascade through individual nodes and connections, and (b) the spread time measure consistently displayed the greatest effect size for the left-right asymmetry (Fig. 1).

To investigate how the cascade spreading trajectories differed between the left and right auditory cortices, we further investigated spread times to specific targets.

Fig. 2 shows spread times for the left and right auditory seeds separately, stratifying the target ROIs into those contralateral and ipsilateral to the auditory seed node. We note three trends: (a) consistent with Fig. 1, spread times are generally faster from the right auditory seed, (b) spread times are faster for ipsilateral compared to contralateral targets (*P* = 5.19 *×* 10*^−^*^6^ and *P* = 1.35*×*10*^−^*^7^ for left and right auditory cortex, respectively), and (c) the biggest discrepancies between ipsilateral and contralateral targets are observed for temporal lobe targets, suggesting that much of the observed asymmetry is driven by less efficient communication between the left auditory cortex and the contralateral temporal lobe. Comparable results were observed in the two replication datasets, where signals originating from right auditory cortex reach the contralateral temporal lobe faster than signals originating from the left auditory cortex (Fig. S1).

**FIG. 2.**
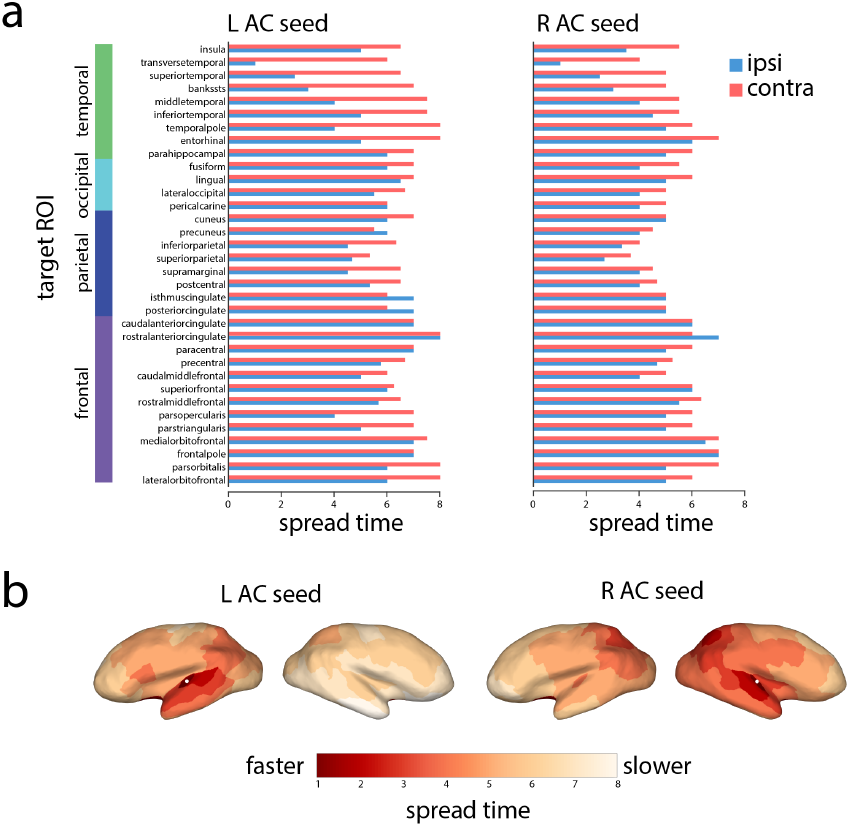
Simulated spreading from auditory cortices to specific target regions. (a) Spreading times to other nodes of the network, separated by lobe and hemisphere (blue for ipsilateral areas, orange for contralateral areas). (b) Spreading times for left and right auditory seeds projected to the cortical surface. The projected locations of the primary auditory nodes are indicated by white dots.

### No comparable asymmetry in the visual system

While our results suggest a consistent asymmetry of the auditory cortices, it is possible that this lateralization is not unique to the auditory system, but perhaps a more general feature of sensory systems. To investigate this possibility, we repeated the analyses described above, but with a focus on primary visual (pericalcarine) cortex (Fig. 3). Unlike the auditory cortex, there was no evidence to suggest hemispheric asymmetry, with no statistically significant differences in path length (*P* = 0.30), communicability (*P* = 0.80) or spread time (*P* = 0.55).

**FIG. 3.**
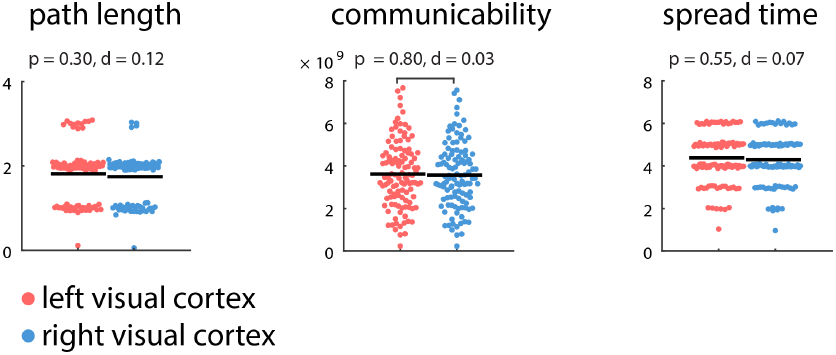
Communication distance from primary visual cortices to the rest of the brain. The centrality of left and right visual cortices was estimated by their topological distance to other brain areas in terms of path length, communicability and spread time. Shorter path length, greater communicability and shorter spread times indicate greater proximity. Mean values for each distribution are indicated by solid horizontal black lines. For visualization, a random horizontal jitter was added to all points. In the case of path length, which is a discrete-valued variable, an additional vertical jitter was added to all points.

## DISCUSSION

The present investigation reveals a network-level asymmetry of the auditory system across multiple datasets. We highlight three principal findings: (1) the right auditory cortex is better integrated in the connectome, facilitating more efficient global communication, (2) these differences emerge only when communication processes are assumed to involve more than just the topologically shortest paths, and (3) much of the asymmetry is driven by differences in communication pathways to the opposite hemisphere.

### Lateralization of auditory networks

These findings support the notion that the functional asymmetry of the auditory system can be at least partly attributed to its embedding in the global anatomical network. Converging evidence from activation studies [27, 67], functional connectivity [13, 48], anatomical connectivity [1, 11] and stimulation [1, 2] points toward the possibility that the right auditory cortex is better positioned to communicate with and influence other systems. Our results suggest that the asymmetry could be at least partly explained by differences in several anatomical pathways, and that these differences potentially “accumulate” as information travels from auditory cortex towards more distant areas.

Specifically, our results suggest that the asymmetric influence of left and right auditory cortex is most pronounced with respect to the contralateral hemisphere. Spreading toward proximal areas in the ipsilateral hemisphere proceeded at a comparable pace for the two seeds, but the differences became more pronounced as the cascades coursed through the contralateral hemisphere, with the greatest differences observed for contralateral temporal lobe areas (Fig. 2). Building on previous reports that individual differences in the strength of auditory transcallosal pathways are related to TMS-induced modulation of interhemispheric functional connectivity [1], our results point to the possibility that these asymmetries are also partly due to indirect pathways. Interestingly, a previous study of left-right asymmetries found that the connectivity patterns and lateralization of the auditory cortex may be more nuanced; while several anterior-posterior projections emanating from the auditory cortex were stronger on the right, other, mainly ventral-dorsal projections, were found to be stronger on the left [11]. Altogether, these studies raise the possibility that differences in anatomical connectivity impart a distinct functional profile on the left and right auditory cortices.

While the present results emphasize the existence of a lateralized auditory network, it is important to note that this lateralization may be more general and may manifest in other systems as well [16]. Although we found no evidence to suggest a similar lateralization in primary visual cortex, other studies have reported differences in anatomical connectivity for the two hemispheres. For instance, lateralization of connectivity and centrality is observed in several areas, with the right hemisphere displaying a more highly interconnected architecture, resulting in shorter average path lengths [36]. This observation suggests that the rightward asymmetry of the auditory system observed in the present study may be part of a broader pattern and warrants further investigation.

The present findings may be interpreted in the light of long-standing models of hemispheric specialization that have proposed various organizational principles to explain the phenomenon. Half a century ago, based on human lesion data, [55] postulated that hemispheric functional asymmetries could be explained on the basis of more focal representation of function on the left compared to a more diffuse representation on the right. This idea and other related concepts have been debated over many years without resolution [9], one of the problems being the rather vague nature of the description, and the lack of clear neuroanatomical basis. This idea may be reinterpreted in light of the asymmetric patterns of spread of activity: rather than reflecting more diffuse organization as such, the enhanced connectivity of the right auditory cortex with other parts of the brain may lead to greater integration of functional processes across widely distributed areas, which might manifest as a more diffuse pattern in response to lesions; conversely the reduced connectivity of left auditory cortex would be associated with more specific interactions especially within the left hemisphere, leading to more focal lesion effects.

The present findings are also compatible with a longstanding conjecture that hemispheric specialization may be related to interhemispheric conduction times [53], so that computations that require relatively rapid interactions across regions may be better supported by local circuitry within a hemisphere. Speech processing, for instance, has been proposed to depend on critical intrahemispheric computations relating auditory, motor, and other structures within the left hemisphere [48], and may be linked to enhanced auditory temporal resolution [66]. Such speech-related processes may thus benefit from the more focal left-side intrahemispheric organization we describe. A number of prior studies have also reported enhanced white-matter tracts within the left-hemisphere speech system compared to the right [12, 21, 51], in keeping with a more tightly organized intrahemispheric system. As well, Iturria-Medina et al [36] point out that the right hemisphere show higher graph-theoretic indices of efficiency and interconnectivity than the left, again broadly consistent with our findings. Other, more local patterns of anatomical asymmetries within auditory cortices that have been described in the literature [35, 42, 43, 52] can now also be re-examined in light of the long-range anatomical connectivity asymmetries described here.

Our anatomical findings also fit well with more recent reports of functional asymmetries. For example Tomasi and Volkow [58] report greater short and long-range connectivity in the right temporal cortex compared to the left, consistent with better transfer of information from right auditory-related areas to the rest of the brain. Liu et al. [41] report greater left-hemisphere functional connectivity from resting-state date for several seed regions, including left superior temporal gyrus, indicating that this region has greater exchange of information with other left-hemisphere structures than its homologue on the right. Similarly, Gotts et al [29] report that resting-state connectivity patterns support greater within-hemisphere interactions on the left side (segregation) but greater between-hemisphere interactions on the right side (integration), a pattern that is consistent with our observations. Importantly, these authors demonstrated that the degree of integration vs segregation is related to individual differences in performance on cognitive task, thus demonstrating that degree of lateralization is related to behavioral ability (but see [12]), and raising the possibility that cognitive function or dysfunction may in future be linked to the patterns of communication uncovered with the present methods.

### Beyond shortest path communication

Interestingly, this auditory asymmetry is not observed when communication is assumed to occur exclusively along shortest paths, and only emerges when additional communication pathways are taken into account, as operationalized by the communicability and spread time measures. These results are part of a growing realization that distributed communication and synchronization in brain networks may proceed via alternative routes [5, 26, 30], with several recent methods developed to quantify path ensembles [3, 4, 17, 18, 25, 32] and the potential for pathways to participate in the diffusion of information [28, 46, 47]. Other recent methodologies revolve around similar ideas, including controllability of linear time-shift invariant systems [7, 34], activity flow mapping [15, 33] and simulated perturbations of ongoing oscillatory dynamics [14, 56]. Our results highlight the need to consider the form of communication that a particular measure assumes [8], and that measures founded exclusively on the concept of shortest paths may not adequately capture the richness and complexity of distributed computations in brain networks [5, 26].

### Methodological considerations

The replicability of the effect suggests robust asymmetries in the auditory system, but it is important to note several limitations as well. First, all our conclusions are based on networks reconstructed from diffusion weighted imaging, a method known to be susceptible to false positives and negatives [37, 57]. Although we attempted to mitigate inaccuracies that may be present at the single-subject level by focusing on group-consensus networks derived from high-quality acquisitions (e.g. DSI, HARDI) in large samples of participants, and by repeating our analyses in multiple datasets, systematic errors or biases in the tractography procedure may still be present. In addition, networks derived from diffusion imaging are by definition undirected, limiting inferences about directionality of influence. These considerations highlight the need for new technologies for noninvasive mapping of white matter projections in the human brain.

Second, our ability to capture network asymmetries is contingent on the accuracy of our communication models. All network measures âĂŞ including simple path length âĂŞ assume some form of communication [8], but how information is transferred among topologically distant neural elements remains unknown. Although we estimated the centrality of the auditory network across a spectrum of communication mechanisms, from shortest path communication to diffusive spreading, it is nevertheless possible that inter-regional communication proceeds via a different mechanism.

### Conclusion

The present study highlights how the network configuration and embedding of a particular region may contribute to its functional lateralization. As our ability to image, reconstruct and stimulate specific neural circuits advances, theoretical models of how perturbations and influence spread through brain networks will become increasingly important. These techniques will ultimately help to create a closer correspondence between structural and functional properties of specific areas and systems.

## ACKNOWLEDGMENTS

BM acknowledges support from the Natural Sciences and Engineering Research Council of Canada (NSERC Discovery Grant RGPIN #017-04265) and from the Fonds de recherche du Québec – Santé (Chercheur Boursier). RZ is supported by funding from the Canadian Institutes of Health Research and the Canada Fund for Innovation. XNZ was supported by grants from the National Basic Research (973) Program (2015CB351702), the Natural Science Foundation of China (NSFC 81471740), Beijing Municipal Science and Tech Commission (Z161100002616023), and the Major Project of National Social Science Foundation of China (14ZDB161). XNZ and OS are members of an international collaboration team supported by the NSFC Major Joint Fund for International Cooperation and Exchange (81220108014).

---

**FIG. S1.**
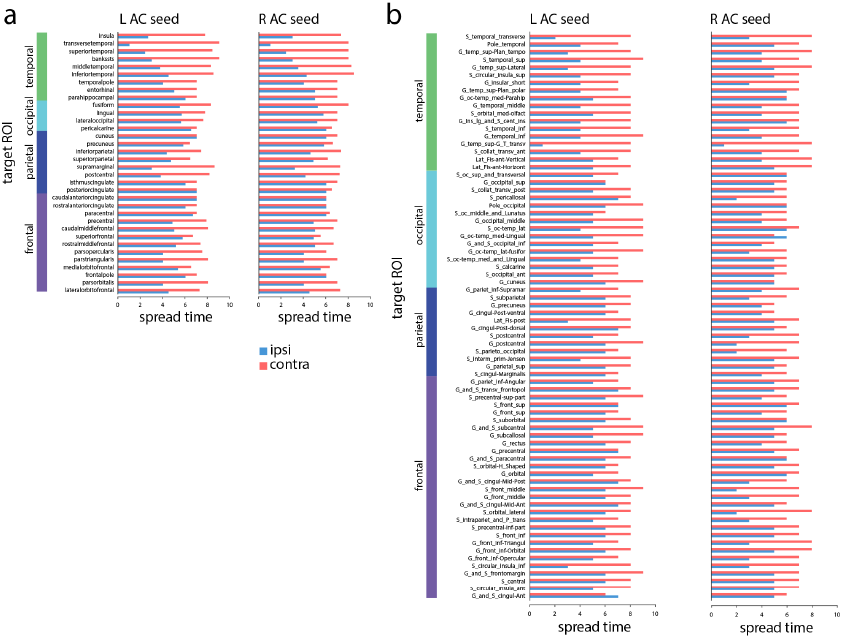
Simulated spreading from auditory cortices to specific target regions in the (a) HCP and (b) NKI datasets. Perturbations are introduced in the left and right auditory cortices (L and R AC). Spreading times to other nodes of the network are stratified by lobe and hemisphere (blue for ipsilateral areas, orange for contralateral areas).

**FIG. S2.**
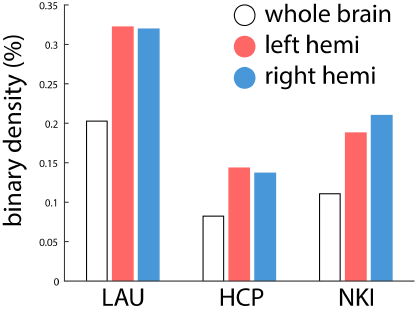
Binary density of the group-level networks. Density is shown for each group-consensus network, as well as for each of the subgraphs corresponding to the hemispheres.

**FIG. S3.**
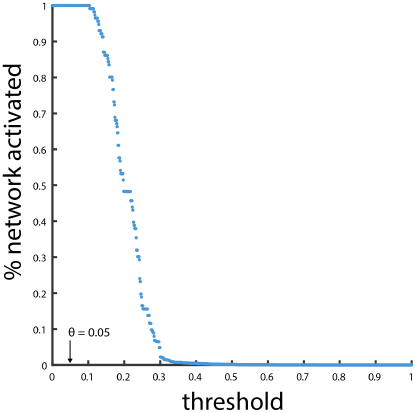
The effect of threshold on global spreading. The linear threshold model (LTM) was used to simulate the effects of focal perturbations at each of the 114 nodes in the Lausanne (LAU) dataset. At low thresholds, the whole network can be trivially activated. As the threshold is increased, some nodes become increasingly difficult to activate, and the total proportion of the network that is activated begins to decrease.

**FIG. S4.**
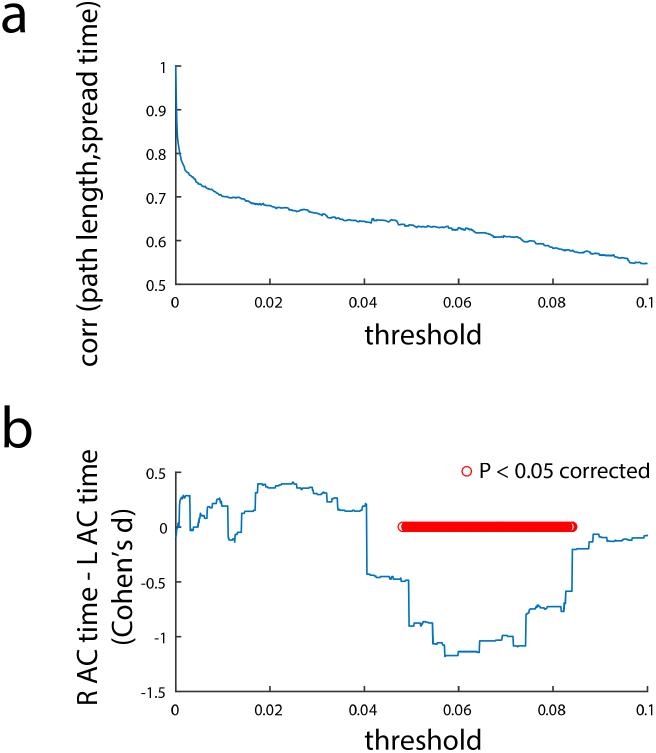
Choosing a threshold. (a) The effect of threshold on the correlation between spread time and path length. When the threshold is low, the dynamics of the model resemble a breadth-first search and spread times are perfectly correlated with path length. As the threshold is increased, spreading is driven away from shortest paths (as evidenced by the decreasing correlation) and more akin to diffusion. (b) The effect of threshold on left-right auditory cortex (AC) asymmetry. The difference in spread time between left (L) and right (R) auditory cortex is shown as a function of threshold. The difference is quantified as a Cohen’s d effect size. At lower thresholds, when spreading is similar to shortest path routing, there are no significant differences between L and R AC. As the threshold is increased, there is a range in parameter space where spreading is significantly faster from the right auditory cortex compared to left auditory cortex (Wicoxon rank sum test, FDR corrected).

